# PRIME: a probabilistic imputation method to reduce dropout effects in single cell RNA sequencing

**DOI:** 10.1101/2020.01.03.893867

**Authors:** Hyundoo Jeong, Zhandong Liu

**Affiliations:** Department of Mechatronics Engineering, Incheon National University, Incheon, 22012, R.O.Korea; Jan and Dan Duncan Neurological Research Institute, Texas Children’s Hospital, Houston, TX 77030, USA; Department of Pediatrics, Baylor College of Medicine, Houston, TX 77030, USA

## Abstract

Single-cell RNA sequencing technology provides a novel means to analyze the transcriptomic profiles of individual cells. The technique is vulnerable, however, to a type of noise called dropout effects, which lead to zero-inflated distributions in the transcriptome profile and reduce the reliability of the results. Single-cell RNA sequencing data therefore need to be carefully processed before in-depth analysis. Here we describe a novel imputation method that reduces dropout effects in single-cell sequencing. We construct a cell correspondence network and adjust gene expression estimates based on transcriptome profiles for the local community of cells of the same type. We comprehensively evaluated this method, called PRIME (**PR**obabilistic **IM**putation to reduce dropout effects in **E**xpression profiles of single cell sequencing), on six datasets and verified that it improves the quality of visualization and accuracy of clustering analysis and can discover gene expression patterns hidden by noise.

## Introduction

The rapid development of single-cell RNA sequencing technologies (1; 2; 3; 4) have enabled researchers to acquire detailed transcriptomic profiles for individual cells in a high-throughput manner. This technology provides an important means for studying cell-to-cell variability, and it is becoming a critical tool for a variety of research endeavors, including cell type identification, pseudo time ordering, and deconvolution of heterogeneous samples (5; 6; 7; 8; 9).

The principle drawback of current single-cell RNA sequencing technology is its vulnerability to technical and biological noise. Individual cells have only a very small amount of mRNA (compared to tissue samples), which requires enormous amplification before analysis. A low initial quantity of a particular transcript can mean that it will be completely missed during the reverse transcription and DNA amplification steps, and thus will not be detectable by subsequent sequencing. Neighboring cells can have wide variability in gene expression, such that a gene expressed at a moderate or high level in one cell is expressed at a low level in another and thus fails to be detected, leading to a ‘false zero’ known as a dropout event. Single-cell RNA seq data is notorious for producing an excessive number of artificial zeros in the expression profile, which must be distinguished from ‘true zeroes.’ Several new analytic tools have been developed using zero-inflated models (10; 11; 12), but most of the current genomic tools were developed based on the distribution of bulk sequencing data (13; 14), which is inappropriate to the nature of single-cell sequencing data.

Several computational methods have been developed to reduce the dropout events by imputing the missing values in single cell sequencing (15; 16; 17; 18). SAVER (15) models single cell gene expression with UMI (unique molecular identifier) counts through Poisson-Gamma mixture and estimates the prior parameter using Poisson lasso regression. Then it recovers the dropouts based on the weighted average of the observed and predicted counts. DrImpute (17) identifies the set of similar cells through k-means clustering and imputes the missing values by averaging the gene expression in the same cluster. To enhance the robustness of the imputation results, DrImpute averages them for multiple k parameters. ScImpute (16) estimates the dropout probability through a mixture model, where it models the gene expression as a Gaussian distribution and the zero-inflated dropout event as a Gamma distribution. It then imputes only those genes with a high dropout probability by utilizing gene expression values from similar cells that are less affected by the dropout events. MAGIC (18) constructs a Markov transition matrix to represent similarities between cells and powers the matrix up to t times in order to model a heat diffusion process. Then, it imputes the missing values through the weighted average of the same genes for the neighboring cells in the Markov affinity matrix.

In this paper, we propose a novel imputation method, called **PRIME**(**PR**obabilistic **IM**putation to reduce dropout effects in **E**xpression profiles of single cell sequencing), to effectively deal with dropout events in single-cell RNA sequencing. First, we construct a cell correspondence network through similarity measurements across cells (Figure 1). Next, we identify the local community for a target cell by using an efficient random walk protocol. Finally, we impute the gene expression in the target cell based on the probabilistic weight parameter, which is computed based on the variance of gene expression in the local community. We perform these steps until it meets the stop conditions (Figure 1).

**Figure 1.**
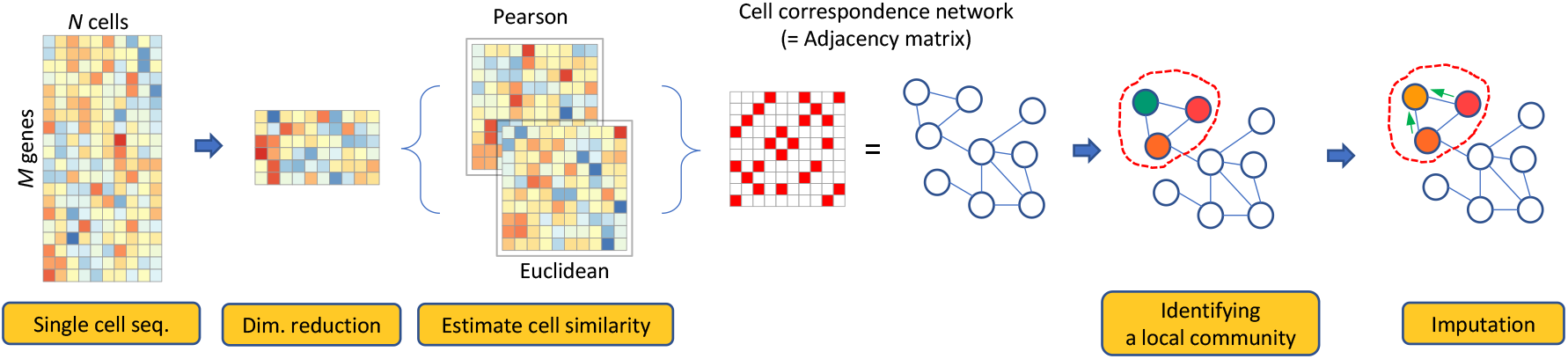
Overall workflow of PRIME. PRIME reduces the dimensionality of the datasets and estimates two similarities matrix using Euclidean and Pearson correlation. Next, it constructs a network by inserting edges between similar cells. Finally, PRIME adjusts expression levels using the average expression levels of its neighboring cells. This process is iterated till convergence.

## Materials and Methods

### Datasets and preprocessing

To assess and compare the performance of single-cell imputation methods, we utilized six single-cell RNA sequencing datasets. i) Buettner et al. (19) provide single-cell RNA sequencing datasets for mouse embryonic stem cells at different cell cycle stages. There are 59, 58, and 65 cells in G1, G2M, and S phase, respectively. The read count for the cell cycle genes is provided in the supplementary file in (19). ii) Usoskin et al. (20) provided single-cell RNA sequencing data for mouse sensory neurons (for peptidergic nociceptors (PEP), non-peptidergic nociceptors (NP), neurofilament containing (NF), and tyrosine hydroxylase containing (TH)). The raw sequencing data is available at gene expression omnibus (GEO) with accession number GSE59739. iii) Zeisel et al. (21) performed large-scale single-cell RNA sequencing on the mouse somatosensory cortex and hippocampal CA1 region. In this dataset, there are seven major cell types that can be classified into 47 different subclasses. We retained only the four major cell types (interneurons, oligodendrocytes, pyramidal CA1, and pyramidal SS neurons). The raw data is archived at the GEO with the accession number GSE60361. iv) The Darmanis dataset (22) provided single-cell RNA sequencing for human brain, and we removed only the cell type labeled ‘hybrid’. The raw data is deposited at GEO with the accession number GSE67835. v) Chu et al. (23) provided bulk and single- cell sequencing for human embryonic stem cells and also the time series sequencing for cell differentiation to endoderm. The raw data is archived at GEO with the accession number GSE75748. For all datasets, we remove genes that are not expressed across all cells. The number of cells and cell types for different datasets are summarized in Table 1.

**Table 1.** Single cell RNA sequencing datasets.

| Data source | # cells | # cell types | Source |
| --- | --- | --- | --- |
| Buettner et al. | 182 | 3 | Mouse embryonic stem cells |
| Usoskin et al. | 622 | 4 | Mouse sensory neurons |
| Zeisel et al. | 2,448 | 4 | Mouse brain |
| Darmanis et al. | 366 | 4 | Human brain |
| Chu et al. | 1,018 | 7 | Human embryonic stem cells |
| Chu et al. (Chu.time) | 758 | 6 | Human definitive endoderm cells |

### Parameter settings for each algorithm

To compare the performance of PRIME with SAVER (15), scImpute (16), DrImpute (17), and MAGIC (18), we utilized R implementation to run each method with the respective default parameters. If the method supports parallel processing, we utilized the maximum number of CPU cores. In this study, we utilized 4 CPU cores for SAVER and scImpute. We tested scImpute based on the default parameters and set the number of the clusters as the number of cell types. For a fair comparison, we tested scImputed without the true label for each cell type, as the other methods do not require the true label. To run DrImpute, we utilized the default parameters and performed the normalization as recommended in the package. We also utilized the default parameter for MAGIC so that it optimizes the diffusion parameter based on their own criterion.

### Data normalization and network construction

The proposed single-cell imputation method consists of three major steps: i) constructing a cell correspondence network, ii) identifying a local community for each cell, and iii) performing a probabilistic imputation for each gene expression. These three steps continue until the maximum number of iterations has been reached or there are no meaningful changes in the imputed expression values compared to the previous iteration. The basic intuition of the iterative approach is that since the raw single-cell RNA sequencing data could be corrupted by technical noise such as dropout events, it is unreliable to impute the dropouts based on noisy datasets. As we impute the technical noise, the reliability of the dataset increases, which improves imputation results in the next iteration.

To begin, suppose that we have single-cell RNA sequencing data and it can be represented as an *M* by *N* dimensional matrix, where *M* is the number of genes and *N* is the number of cells. We normalize the library size of the single-cell RNA sequencing data matrix using a cpm (counts per million) and take a log-transformation. We have a normalized input **X**_**N**_, which is given by

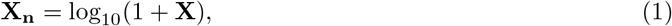

where **X** is cpm transformed input data. Note that the input value is not limited to read counts and it is acceptable if it represents relative expressions of genes across cells. Once we have a normalized data matrix **X_n_**, we start the iterative imputation process by constructing the cell correspondence network based on the cell-to-cell similarity. The cell correspondence network can be represented as a graph 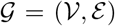, where a node 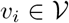 represents an *i*-th cell and the cell correspondence can be represented as an edge 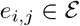 so that if cells *v*_*i*_ and *v*_*j*_ are similar to each other, the edge weight *e*_*i,j*_ can have a positive value. In this study, we utilized both Euclidean distance and Pearson correlation to estimate cell-to-cell similarity. Before estimating the similarity, since a single-cell sequencing generally includes a number of cells and genes, we first reduce the dimension of the input data **X_n_** in order to reduce the computational complexity and shorten the running time of the method. To this end, we select highly variable genes across all cells using Seurat (24) and obtain a low-dimensional representation for each cell using PCA (principal component analysis).

Next, we construct the cell correspondence network (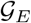) based on the Euclidean distance and the network (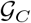) for the Pearson correlation and combine them to obtain a comprehensive cell correspondence network 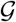. First, we compute the Pearson correlation between each cell using top 20 principal components (PCs) to construct a cell correspondence network for the Pearson correlation. For a given cell *v*_*i*_, we select the cells having a high correlation and consider the cells as the neighboring nodes in the cell correspondence network, 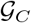. Note that the neighboring nodes indicate the set of cells that can be classified as the same cell type with similar expression patterns, and the neighboring nodes for the cell *v*_*i*_ can be selected based on the following criterion:

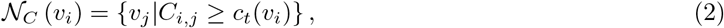

where *C*_*i,j*_ is a Pearson correlation between the cell *v*_*i*_ and *v*_*j*_, and *c*_*t*_(*v*_*i*_) is the threshold to select the neighboring cells. We adaptively select the threshold by taking min {0.85, (90 percentile of *C*_*i,j*_, ∀*j*)}. Then, we insert edges between the cell *v*_*i*_ and the cell 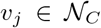. The adjacency matrix for a cell correspondence network based on Pearson correlation is given by

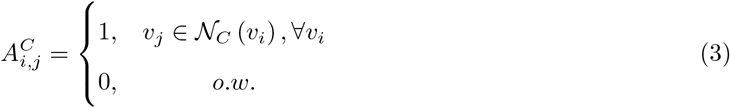

The above adjacency matrix is asymmetric and it generates a directed network because the selection of cell *v*_j_ as the neighbor of cell *v*_*i*_ does not necessarily guarantee the opposite case even though the Pearson correlation matrix is symmetric. To make it an undirected network and give more confidence to the bidirectional edges (i.e., the cell *v*_*i*_ selects the cell *v*_*j*_ as its neighboring node, and vice versa), the adjacency matrix for the undirected network can be obtained by linear combination of **A**_C_ and its transpose, which is given by

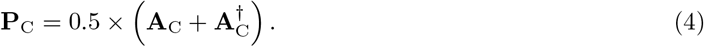

Similar to the cell correspondence network based on the Pearson correlation, we first compute the Euclidean distance between the first 20 PCs for each cell. Since the Euclidean distance becomes smaller as the cell-to-cell similarity increases, we utilize the Gaussian kernel to obtain the Euclidean similarity, which is given by

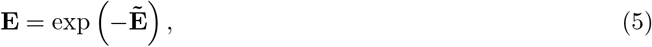

where 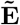 is the element-wise square of the rescaled Euclidean distance matrix 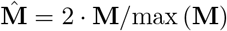 and M is the *|N | × |N |* dimensional matrix representing Euclidean distance between each cell. Then, we can obtain the adjacency matrix **P**_E_ for the cell correspondence network, 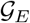, through the same process described in Eq. (2) to Eq. (4).

After computing the adjacency matrices for the Pearson correlation and Euclidean similarity, we combine both similarities through a linear combination with an equal weight and take an element-wise square, where it gives more weight on the edges consistently identified by both metrics and decreases the weight on the edges identified by only one similarity measurement. The resulting adjacency matrix for the cell correspondence network 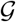 is given by

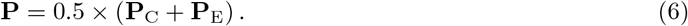

In the network construction step, starting from a directed network with an equal weight, we iteratively rescale the edge weights by performing a linear combination of matrices and element-wise square, and this process gives more weight to the consensus edges (e.g., the edge connecting the cells *v*_*i*_ and *v*_*j*_ identified simultaneously by both Pearson correlation and Euclidean criteria).

### Identifying local community and probabilistic imputation

To reduce the dropout effects in single-cell sequencing in a biologically unbiased manner, it is necessary to distinguish dropouts from true biological zeros (16). To this end, we take advantage of the surrounding cells. The basic intuition is that if the gene is not expressed in the majority of cells of the same type, the observed zero is highly likely to be a true zero. However, if the gene has a positive expression in the most of the cells in the same type (i.e., cells in the local community) but it is not expressed in a particular cell, the detected zero has a higher probability of being a dropout event, where it should be recovered to the true (or expected) values. Thus, we compare the gene’s expression in a particular cell to its expected expression in the set of similar cells – a local community identified by a random walk approach inspired by local graph partitioning using a personalized PageRank (PPR) vector (25). Although the PPR vector can be used to identify the exact local network clustering, it has high computational complexity when dealing with the large-scale networks. Since our aim is not to derive the whole set of cells of the same type but to identify a reasonable local community, we adopt a heuristic approach to approximate the PPR vector. First, we perform a column-wise normalization in order to obtain the legitimate stochastic matrix (i.e., the transition probability matrix for the random walker). Then, we obtain the transition probability matrix for the random walker over the cell correspondence network 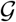 by performing a matrix product to consider a secondary structural similarity.

Next, to identify the set of similar cells for the cell *v*_*i*_, we identify the local community in the cell correspondence network 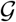 through a random walk. Starting from the *i*-th cell *v*_*i*_, the random walker performs a random movement for *J* steps over the cell correspondence network 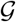 and identifies the local community for the cell *v*_*i*_ by selecting *K* neighboring cells based on the visiting frequency of the random walker for each node. Note that the parameter *J* is empirically set to 3 and *K* is selected by 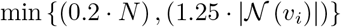,, where 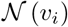 is the neighboring nodes for the cell *v*_*i*_ in 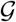 and *N* is the number of cells. After identifying a local community for the cell *v*_*i*_, we computed the mean vector *μ*_*i*_ and variance vector *σ*_*i*_ for *M* genes in the local community, where the *m*-th element in these vectors is the mean and variance for the *m*-th gene of the cells in the local community. Then, we impute the expression *x*_*i*_ for *M* genes in the cell *v*_*i*_ based on the mean and variance of the expression values in the local community. That is, if the *m*-th gene expression in the cell *v*_*i*_ is reliable (i.e., similar to the mean value for the local community), we will assign more weight to the expression of cell *v*_*i*_ and utilize the least information from the neighboring cells to adjust the gene expression values. However, if it significantly deviates from the local mean toward zero, we will assign more confidence to the information obtained from the neighboring cells so that the potential dropout events can be recovered. Based on this intuition, the updated rule for gene expression in the cell *v*_*i*_ is given by

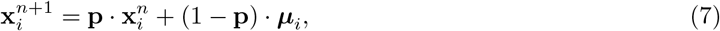

where **p** is the probabilistic weight for the current gene expression and the probabilistic weight **p** is determined by the following sigmoid function:

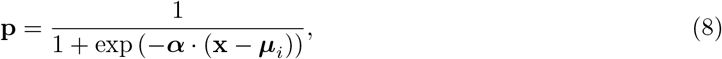

where *α* is scaling coefficients for the sigmoid function and it is selected using the following criterion: min {2 · exp (−0.05 ·*σ*_*i*_), 1}. To avoid the extreme case, we set the marginal value for the scaling coefficient *α* as 1 because if the *α* is set to very low value, the sigmoid function can approximate a step function.

We perform the iterative imputation process until it converges or exceeds the maximum number of iterations. Empirically, we stop the iteration if *|***X**_*n+1*_ *−* **X**_*n*_|^2^*/*(*|M | · |N |*) is smaller than 0.05 or the number of iteration exceeds five.

## Results

### Better identification of cell types from expression profiles

In analyzing single-cell RNA sequencing datasets, the fundamental first step is to visualize each cell in a low-dimensional space and to identify known and novel cell types through clustering-based methods. Dropout events can decrease cell-to-cell similarity within the same cell type, resulting in mistaken identification of cell type. To visualize single cells in a low-dimensional space, we utilized the cell type labels reported in the original papers and employed two popular dimensional reduction methods, PCA and t-SNE (26). Since the imputation recovers dropouts in each cell and depends strongly on the frequency of the dropout events for each cell, the total number of counts for each cell can change dramatically after the imputation. We therefore re-normalized each single cell using the library size after the imputation. In fact, MAGIC normalizes the imputation result by default, but the other methods do not consider post-normalization after imputation. For a fair comparison, we re-normalized the imputed gene expression matrix using counts per million (cpm) and perform a log-transformation to obtain low-dimensional visualizations. Our approach was able to impute dropout values across different cell types and led to a better separation between different cell types (Figure 2).

**Figure 2.**
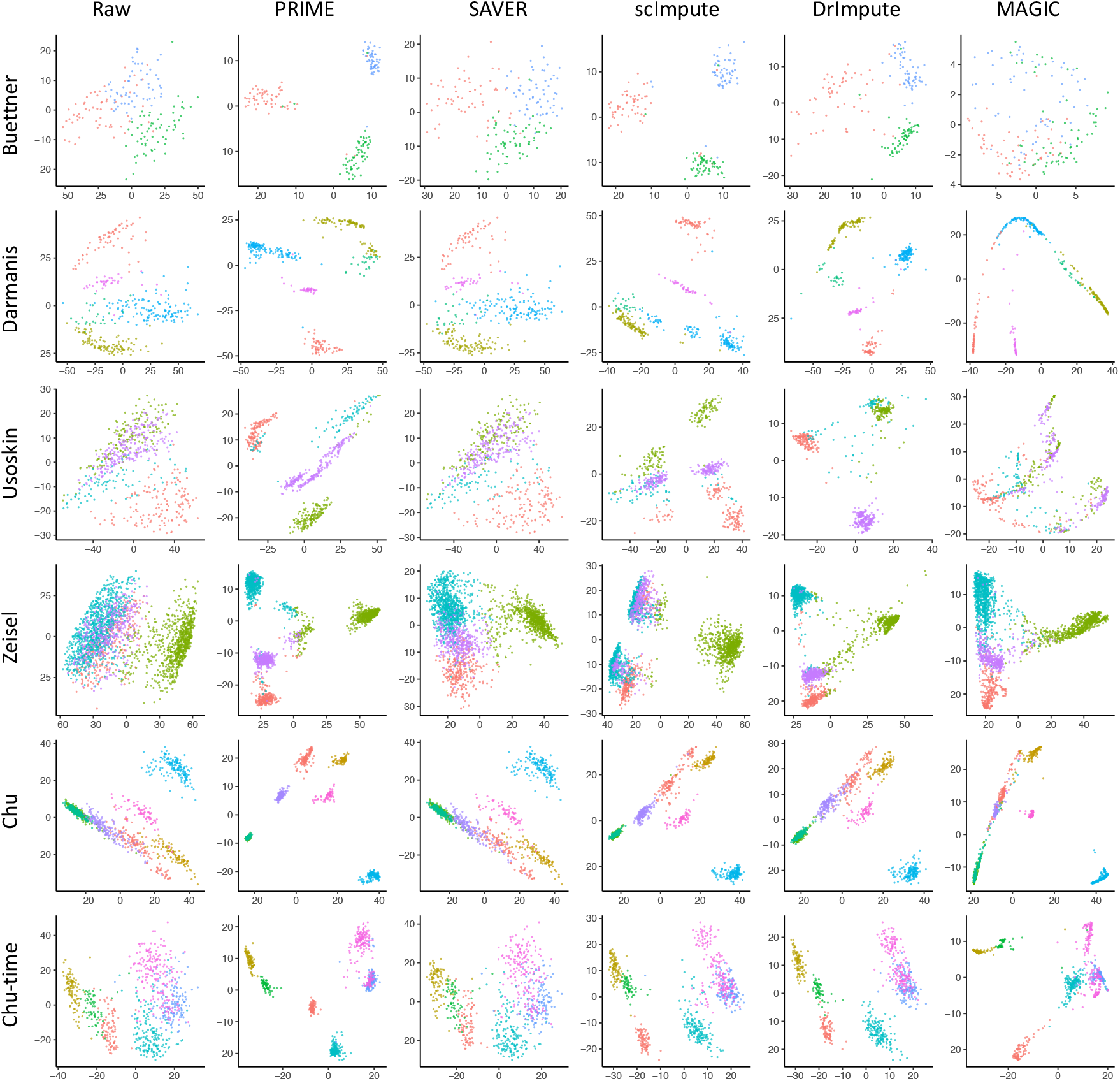
Low dimensional embedding of imputed scRNAseq data over six benchmark datasets. The first two components in the Principal component analysis (PCA) were plotted. PRIME tends to yield a more clear separation for different cell types while preserving the original features demonstrated in the PCA plot using raw input.

We compared our method to four other approaches on six datasets (19; 20; 21; 22; 23) with the cell type labels provided by each original publication. PRIME generated clear visualization results for all six datasets (Figure 2). SAVER showed a negligible effect on the visualization results for most cases. One possible explanation is that SAVER assumed that the gene count can be modeled as a Poisson-Gamma mixture, which can be effective for single-cell sequencing based on UMI counts. But if the count does not follow a Poisson-Gamma mixture distribution, SAVER could fail to effectively impute the dropouts. Even though the count data fits the assumption (e.g., Zeisel dataset (21)), SAVER does not clearly separate different cell types in a low-dimensional space. ScImpute showed a comparable result to PRIME in the Buettner dataset (19), but in the Usoskin (20) and Zeisel (21) datasets, it divided the same cell types into different clusters and merged different cell types into the same cluster. MAGIC failed to separate cell types in the low-dimensional space in most of the datasets. PRIME produced better visualization results than the competing algorithms; the t-SNE plots for the different imputation methods are provided in the Supplementary Figure S1. Overall, PRIME improves the visibility of single cells when compared to the raw datasets.

To measure the improvement on cell type clustering of imputation methods, we used pcaReduce (27), hierarchical clustering based on a normalized Euclidean distance among cells, and spectral clustering using the first two principal components (PCs) as in (16). We obtained spectral clustering through R package, `speccalt` (28). We evaluated the quality of the clustering based on ARI (adjusted rand index), NMI (normalized mutual information), Jaccard index, and Purity. PRIME outperformed the other methods (Supplementary Figure S2). These results demonstrate the effectiveness of PRIME in improving cell type discovery.

### Uncover cell state-dependent gene expression patterns

To demonstrate that effective imputation methods can reduce dropouts and lead to the discovery of hidden gene expression patterns in the single-cell data, we compared all the methods on a set of transcriptomic profiles (19) of 182 mESCs over different cell cycle stages (G1, G2M and S). Cells belonging to the same cell cycle phase were not clustered together using hierarchical clustering (i.e., the color label for the column annotation). In each cell cycle phase, the expression of cell cycle genes typically changes in a periodic pattern—one not observable at the normalized raw single-cell expression level (Figure 3). To determine whether PRIME can improve the signal-to-noise ratio and detect cell cycling patterns, we identified differentially expressed genes with a large fold-change across different cell cycle stages in the raw dataset using DEseq2 (13) for 892 cell cycle genes reported in (19). Then, we plotted the heatmap for differentially expressed genes with a row-wise normalization to visualize cyclical patterns in gene expression in the different cell cycle stages and performed hierarchical clustering to validate the consistency of gene expression at the same stages.

**Figure 3.**
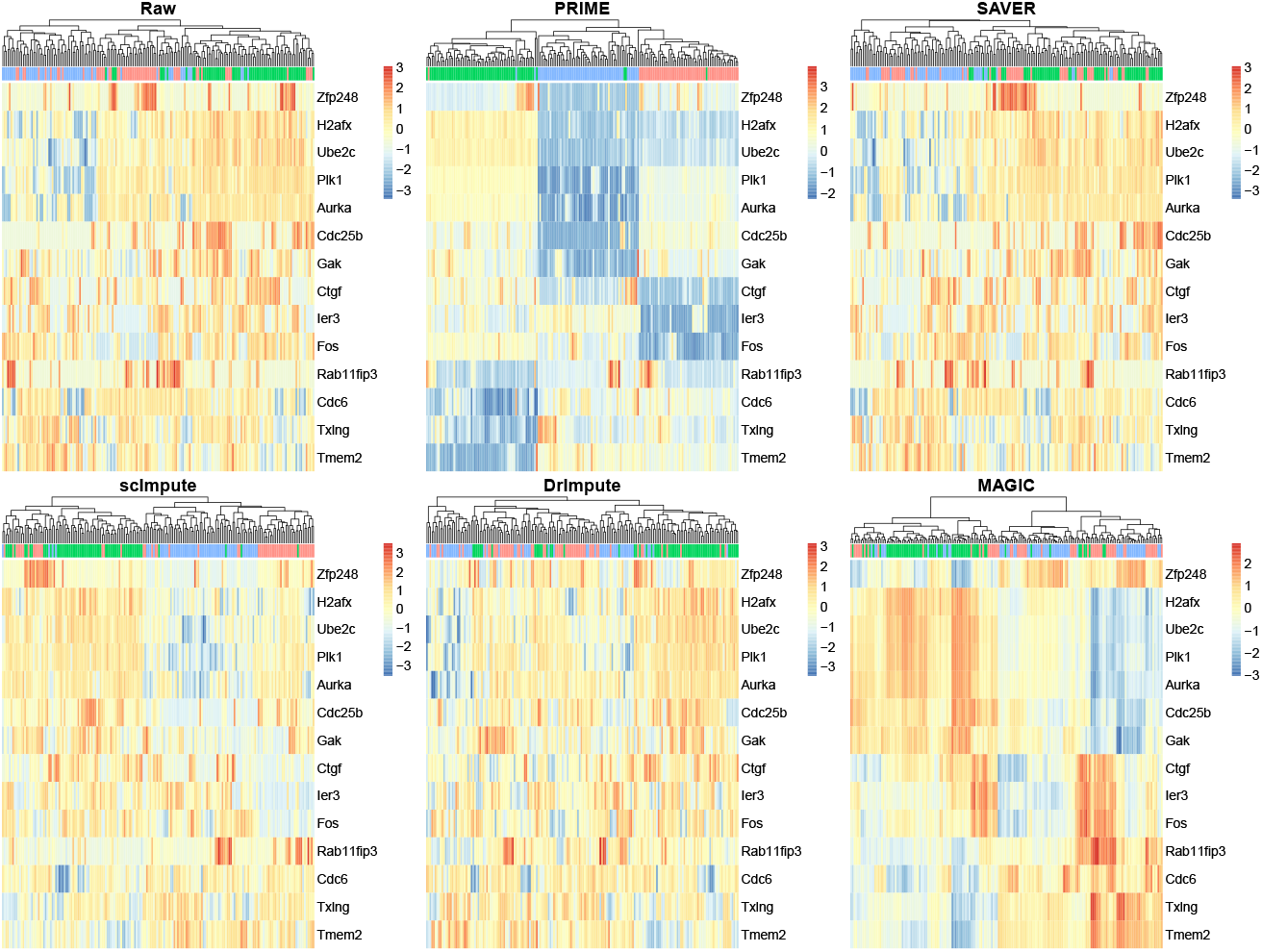
Heatmap plot for the cell cycle genes over single cells at different cell cycle stages. The column color bar indicates the cell cycle stage; blue for G1, red for S phase, and green for G2M. A clear cycling pattern is observed in PRIME imputed data. The similar pattern is only observed MAGIC but not the other impuation methods.

After PRIME imputation, we observed a clear pattern where cells were grouped correctly based on their cell cycle stage. In contrast, no cyclical patterns in DrImpute or SAVER were apparent, and cells across multiple cell cycle phases were clustered together. MAGIC showed a noticeable periodic pattern in the heatmap, but careful examination of the hierarchical clustering results reveals that cells in different phases are clustered together, indicating that gene expression in the same cell cycle phase is highly incoherent. The choice of clustering methods is not the main reason for the mis-clustering of different cell cycle stages (Supplementary Figure S3).

We verified that PRIME recovers dropouts while keeping biological differences between different cycles through the gene expression profiles for the selected genes at different stages of the cell cycle (Supplementary Figure S4). SAVER yields negligible imputation effects and MAGIC decreases biological variations across different cell cycle phases. It clearly shows that the proposed method effectively recovers a greater number of dropouts while maintaining biological heterogeneity across different cycling phases. (Supplementary Figure S5)

### Stable inference on co-expression network

Many bioinformatic studies estimate gene co-expression networks from bulk RNA sequencing data. The number of observations provided by single-cell RNA sequencing —from 500 to 50K cells in a typical experiment—makes this method even more powerful. A large number of dropouts could decrease the stability of the co-expression network, however, resulting in spurious inferences on gene-gene co-expression patterns. To determine whether PRIME can improve the stability of the network inference, we estimated co-expression networks from both raw and imputed data on single-cell profiles generated from human embryonic stem (ES) cells and differentiated definitive endoderm (DE) cells at 0, 12, 24, 36, 72, and 96 hours. We estimated network stability by subsampling 758 cells in this time series dataset. We then used the method of Meinshausen and Buhlmann (29) to estimate over 100 regularization parameters for the network. For a fair comparison, networks estimated from different imputation methods were aligned by their network sparsity level. PRIME consistently identified more reliable edges, no matter the sparsity level (Figure 4). The number of reliable edges for SAVER and DrImpute is smaller than the raw dataset.

For each imputation method, the Meinshausen and Buhlmann method (29) also selected an optimum network at a global stability threshold of 0.95. The optimum network for the raw data has 276 singleton nodes and 391 edges; the optimum network for PRIME has the lowest number of singleton nodes (212) and the largest number of edges (483) of all the methods (Table 2). Since the learned network based on MAGIC is an almost fully connected network that violates the sparsity rule, we excluded it for this comparison.) When comparing the raw data, the proposed method can decrease 64 singleton nodes by identifying more reliable edges, and it can identify 14 percent more reliable edges than the next best algorithm, scImpute.

**Table 2.** Optimum network structures for various methods.

|  | Singleton nodes | Edges |
| --- | --- | --- |
| Raw | 276 | 391 |
| PRIME | 212 | 483 |
| SAVER | 292 | 343 |
| scImpute | 245 | 425 |
| DrImpute | 298 | 332 |

**Figure 4.**
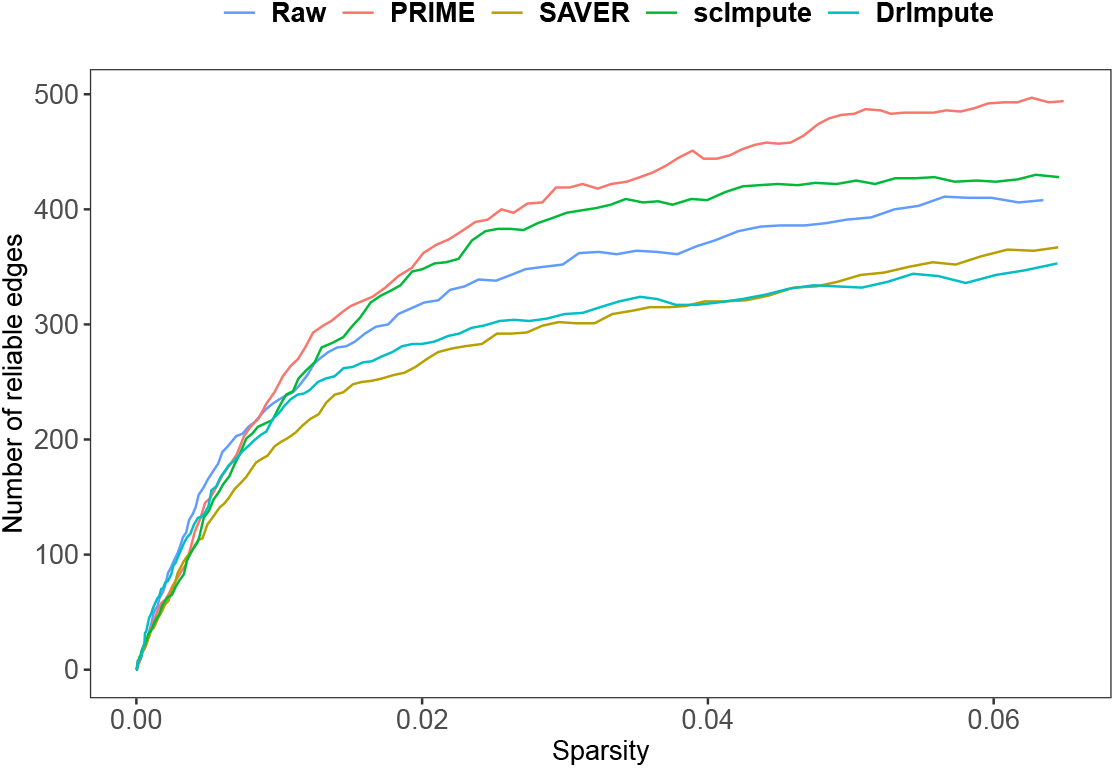
Gene expression network analysis on imputed data. PRIME can detect more edges at a large range of sparsity levels. The number of reliable edges (90% reproducible in sub-sampled data) were plotted against different sparsity levels for all imputation methods.

### Improved correlation with bulk RNA sequencing

Because there tends to be less heterogeneity in single-cell RNAseq data from cell lines than from tissues, single-cell transcriptomes tend to be tightly correlated with bulk RNA sequencing when profiled on cell lines. We took advantage of this property to evaluate the accuracy of various imputation methods. In particular, we used single-cell sequencing for human embryonic stem (ES) cells, where the dataset includes 173 neuronal progenitor cells (NPCs), 138 endoderm derivatives (DE), 105 endothelial cells (ECs), 69 trophoblast-like cells (TBs), 159 human foreskin fibroblasts (HFFs), 212 H1 and 162 H9 human ES cells. For each of the respective cell lines, Chu et al. (23) also profiled the bulk RNA samples at the same time points using Illumina single-end sequencing. Since the gene expression in bulk RNA sequencing approximates the average gene expression of cells in the tissue, bulk sequencing has greater sequencing depth and is less susceptible to dropout events. We therefore hypothesized that effective imputation would increase the ability to find gene expression correlations by effectively removing dropouts.

The correlation between raw data and bulk sequencing was low even though both were sampled from the same cell types. After reducing dropout effects, however, the correlation between bulk and single-cell sequencing clearly increased (Figure 5). MAGIC showed the best performance, and PRIME recorded the second-highest correlation in average. Chu et al. also generate bulk and single-cell RNA sequencing data to produce endoderm derivative cells from human ES cells at different time points (12h, 24h, 36h, 72h, and 96h). Supplementary Figure S7 shows the correlation between bulk and single-cell sequencing at different time points and it shows a similar trend to the Figure 5. Supplementary Figure S8 shows the genes differentially expressed between each pair of adjacent time point at different time points, and it shows that PRIME effectively recovers the gene expression that is close to the bulk sequencing. These results data from single-cell sequencing.

**Figure 5.**
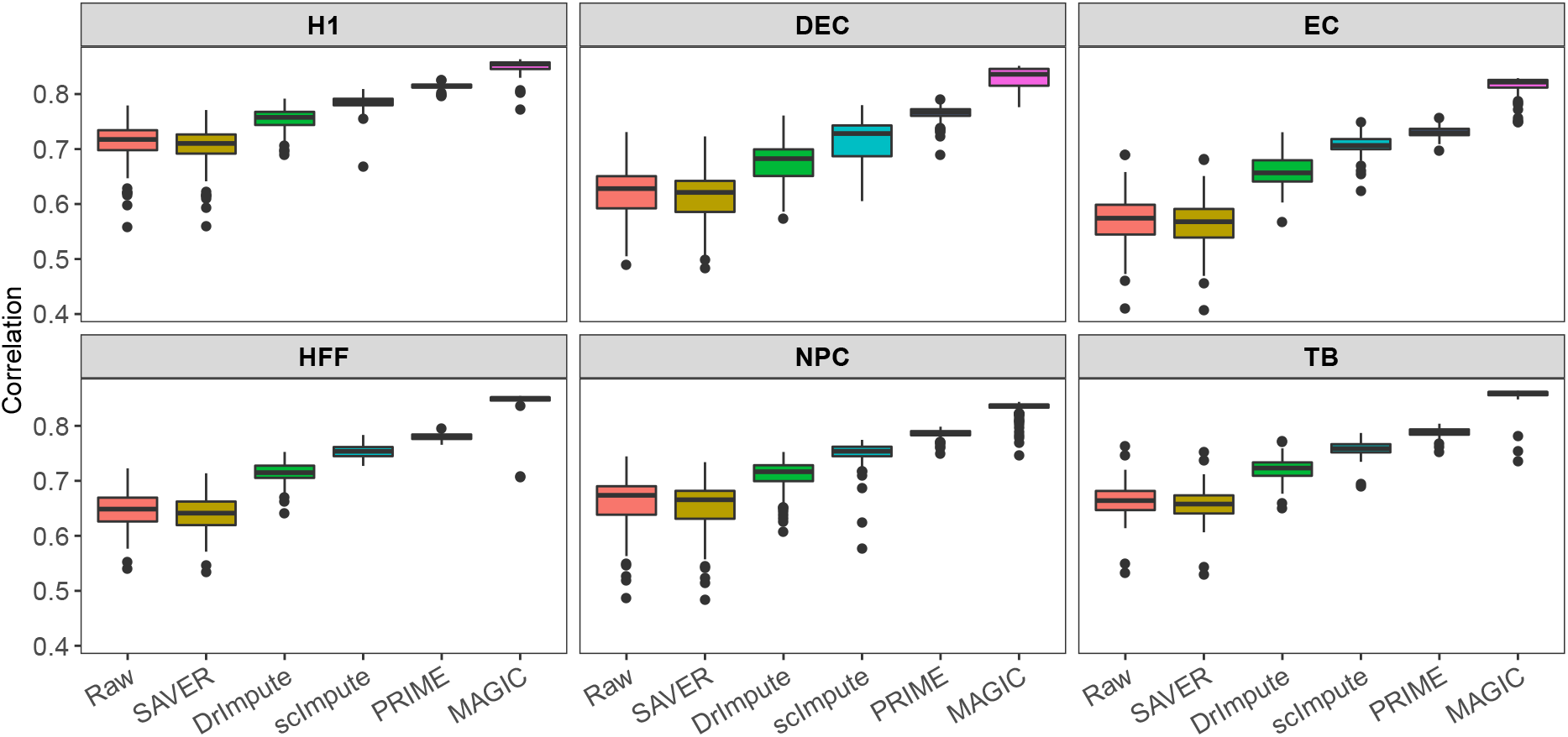
Correlation between scRNAseq and Bulkseq on cell lines. A high correlation is expected between the scRNAseq and Bulk seq. This box plot shows the correlation between bulk expression and the imputed single cell expression using MAGIC and PRIME across multiple cell types.

**Figure 6.**
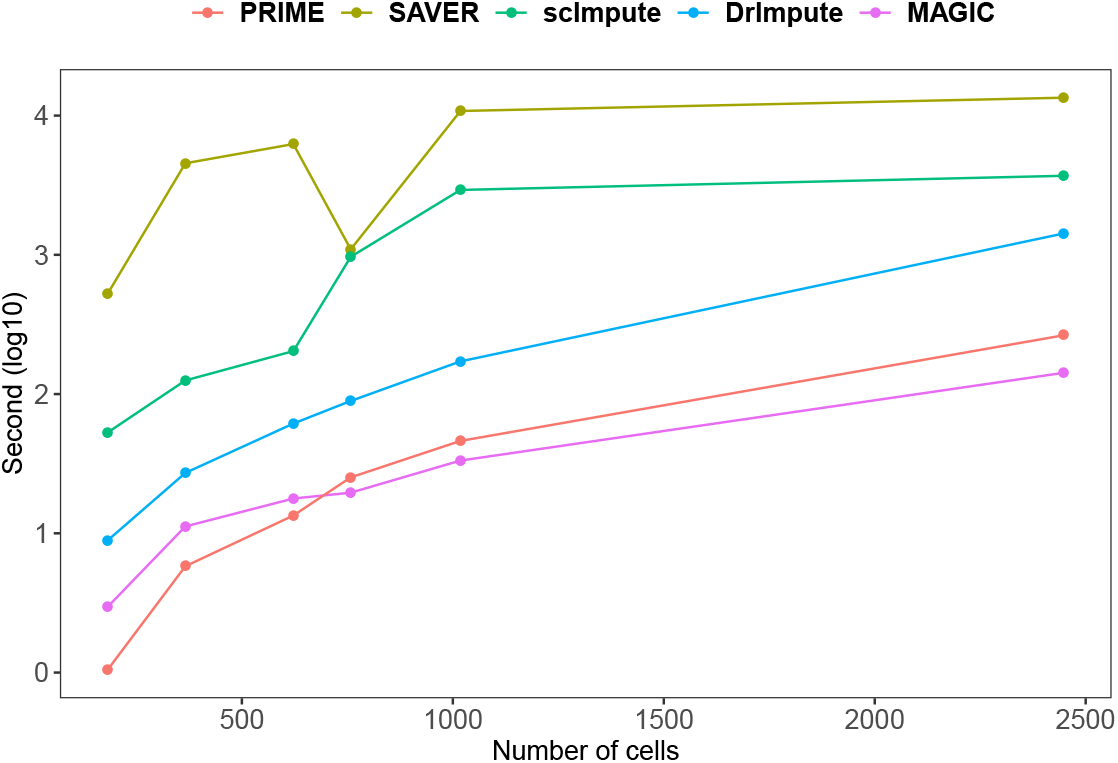
Running time comparison. Six datasets were sorted based on the number of cells sequenced. The running time is plotted on the y-axis in logarithm scale. PRIME and MAGIC are the fastest among these methods. SAVER requires a longer running time compared to all other methods.

### Computation time

One of the major advantages of single-cell RNA sequencing is its ability to profile thousands to millions of cells simultaneously. The scalability of the algorithm is thus an important factor to consider for large-scale single cell analysis. We compared the running time for different imputation algorithms implemented by R script. We utilized SAVER version 1.0, scImpute version 0.0.6, and the latest versions of MAGIC and DrImpute. In this experiment, we used a laptop computer equipped with Intel i5 3.4 GHz and 16 GB RAM. If the algorithm supports a parallelization, we utilized the maximum number of cores (i.e., 4 CPU cores). PRIME and MAGIC required the least computation time even though they utilize only a single core. Although SAVER and scImpute utilized multiple CPU cores, they required much longer computation times and are the least scalable methods.

## Discussion

Here we describe a probabilistic imputation method (PRIME) to reduce dropout effects in a single-cell RNA sequencing data. PRIME iteratively recovers the missing values in single-cell RNA sequencing data based on the expected gene expression in the set of cells that putatively belong to the same cell type. First, we construct a cell correspondence network through Euclidean distance and Pearson correlation, and we identify the local community (i.e., set of highly relevant cells to the imputation target cell) through an efficient random walk protocol to employ the wisdom of crowd. Finally, to decrease dropout events, we impute the gene expressions based on the mean and variance of the gene expressions in the local community. Through a comprehensive evaluation using real-world single cell sequencing datasets, we demonstrate that the proposed imputation method can provide better visualization in a low-dimensional space and cell type clustering, enhanced gene expression patterns, and improved stability for the network inference with rapid computation times. PRIME is also compatible with other single-cell analysis methods. Since it does not change the dimension (i.e., the number of genes and cells) of the input data and it effectively recovers the dropouts in the raw count matrix, it can be directly employed within existing analysis pipelines without complicated and time-consuming manipulation. We propose that this method can be used as the preprocessing step for various single-cell analyses such as a visualization, single-cell clustering, and gene expression analysis. More importantly, the proposed method does not require prior information, such as the number of cell types and cell-type specific marker genes: such information might not be available or it may require additional biological experiments. The proposed method is thus quite practical and versatile to most of the real-world single-cell sequencing studies. It also requires less computational time than the other state-of-the-art algorithms, and it is effective at dealing with large-scale single-cell datasets. Although the proposed method can effectively impute the dropout events in single-cell RNA sequencing, there are certain limitations. For example, if all the genes in a particular cell type are corrupted by dropouts, PRIME would not be able to impute the missing values because there is not enough information from the local community to recover the missing values. In fact, this is a common problem for most of the current imputation methods. To effectively address the problem and improve the accuracy of imputation results, we would integrate an effective data mining strategy with the imputation method. Then, it can automatically identify the prior information such as gene regulatory relationships and cell-type-specific marker genes and accurately infer the missing values even under conditions of extreme dropout events.

## Acknowledgments

This work was also supported by the National Research Foundation of Korea (NRF) grant funded by the Korea government(MSIT) (NRF-2019R1G1A1004803).

